# High Impact: Wikipedia sources and edit history document two decades of the climate change field

**DOI:** 10.1101/2023.11.30.569362

**Authors:** Omer Benjakob, Louise Jouveshomme, Matthieu Collet, Ariane Augustoni, Rona Aviram

## Abstract

Since being founded in 2001, Wikipedia has grown into a trusted source of knowledge online, feeding Google search results and serving as training data for ChatGPT. Understanding the accuracy of its information, the sources behind its articles and their role in the transference of knowledge to the public are becoming increasingly important questions. Meanwhile, climate change has moved to the forefront of scientific and public discourse after years of warnings from the scientific community. Therefore, to understand how it was represented on English Wikipedia, we deployed a mixed-method approach on the article for “Effects of climate change” (ECC), its edit history and references, as well as hundreds of associated articles dealing with climate change in different ways. Using automated tools to scrape data from Wikipedia, we saw new articles were created as climatology-related knowledge grew and permeated into other fields, reflecting a growing body of climate research and growing public interest. Our qualitative textual analysis shows how specific descriptions of climatic phenomena became less hypothetical, reflecting the real-world public debate. The Intergovernmental Panel on Climate Change (IPCC) had a big impact on content and structure, we found using a bibliometric analysis, and what made this possible, we also discovered through a historical analysis, was the impactful work of just a few editors. This research suggests Wikipedia’s articles documented the real-world events around climate change and its wider acceptance - initially a hypothesis that soon became a regretful reality. Overall, our findings highlight the unique role IPCC reports play in making scientific knowledge about climate change actionable to the public, and underscore Wikipedia’s ability to facilitate access to research. This work demonstrates Wikipedia can be researched using both computational and qualitative methods to better understand transference of scientific information to the public and the history of contemporary science.

## 1. Introduction

Wikipedia and its articles feed Google search results, inform Siri and other so-called smart personal assistants, and even help train AI language models like ChatGPT. Launched in 2001, the online encyclopedia is not just a massive database of information but also a trusted source of knowledge in the digital age. Research on Wikipedia shows it generally relies on reliable sources and high-quality academic sources (1–4) even on topics in the public arena like COVID19 (5,6).

From computer sciences (7) to the history of science (8,9), Wikipedia, its articles and the bibliometric data they contain, are increasingly becoming a lucrative and popular research topic. A growing body of research has tried to use Wikipedia for historical purposes, bringing mixed-methods from digital humanities (10), bibliometrics (11,12), textual analysis (13) and others from the disciplines of history and sociology of science and knowledge (14) to the online encyclopedia. These include studies on diverse topics such as the Egyptian revolution (15) and even attempts to map the entire history of human knowledge since the dawn of history (16). Recently, tools have been put forward to consolidate research methods focused on using Wikipedia to study contemporary science; specifically the manner it documents the growth of knowledge and facilitates access to research (9).

Climate change is among the most volatile subjects on Wikipedia (17), and has thus been studied from the perspective of controversy dynamics, in English and Portuguese Wikipedia (17–19). Following years of heated debates that took place mostly in the media, the public consensus around the anthropogenic causes of climate change consolidated, a process that was reflected on Wikipedia, with the controversy appearing to subside from 2010 onwards.

Today it is clear that long-reaching and expanding climatic effects are increasingly a lived reality (20). Climatologists and environmental policy makers have long worked to inform the public about their work (21), and as those become more present the need to access climate knowledge, address denialism, and educate becomes more urgent (22). Climate issues have moved front and center into public discourse (23), with many news outlets now offering climate coverage (24), to name but one example. The United Nations (UN) Intergovernmental Panel on Climate Change (IPCC) has emerged as a key force in addressing climate issues, and consolidating global research efforts on this front (25). From 1990 and until this year, they’ve published numerous reports that laid out the science behind climate change (26) and facilitated global understanding of its ramifications and the threat it poses (27). The IPCC, like the climate change it researched and warned about, was a controversial term on Wikipedia, too; and appeared to also be broadly accepted by 2014, at least in the article for global warming (17).

Given Wikipedia’s unique position online and its ability to bridge academia and the public, and as it also serves as a source of knowledge in its own right, the goal of the present study was to examine how climate change effects were covered on the website. To understand which and how much of the information regarding climate change and its effects was reaching the public through Wikipedia, we used specially developed computational tools to obtain metadata from the website, and combine them with qualitative analyses, including a close comparative reading of past versions of articles. We found that the Wikipedia article in English for the “Effects of Climate Change” (ECC), as well as hundreds of related articles, documented different aspects of the discourse around this topic for over 20 years.

## 2. Bodies of knowledge: Corpus analyses

### 2.1 Corpus delineation

The first stage of our research was defining the body of Wikipedia articles that deal with our topic of interest: climate change and its effects. For this aim, a “corpus builder” (9,28) tool was deployed to scrape Wikipedia’s massive body of English-language articles only for those containing the term “Effects of climate change” in their title or in that of one of their sections. This filtration was aimed at restricting the corpus to articles with significant content relating to climate change effects, rather than those with merely anecdotal mentions, as would appear in the free search directly. This yielded 921 articles which were further divided into sub-corpora: first, those with the term in their main title (“title corpus”), and secondly those that included it in a section’s title (“section corpus”), (Fig 1A and Table S1).

**Fig 1:**
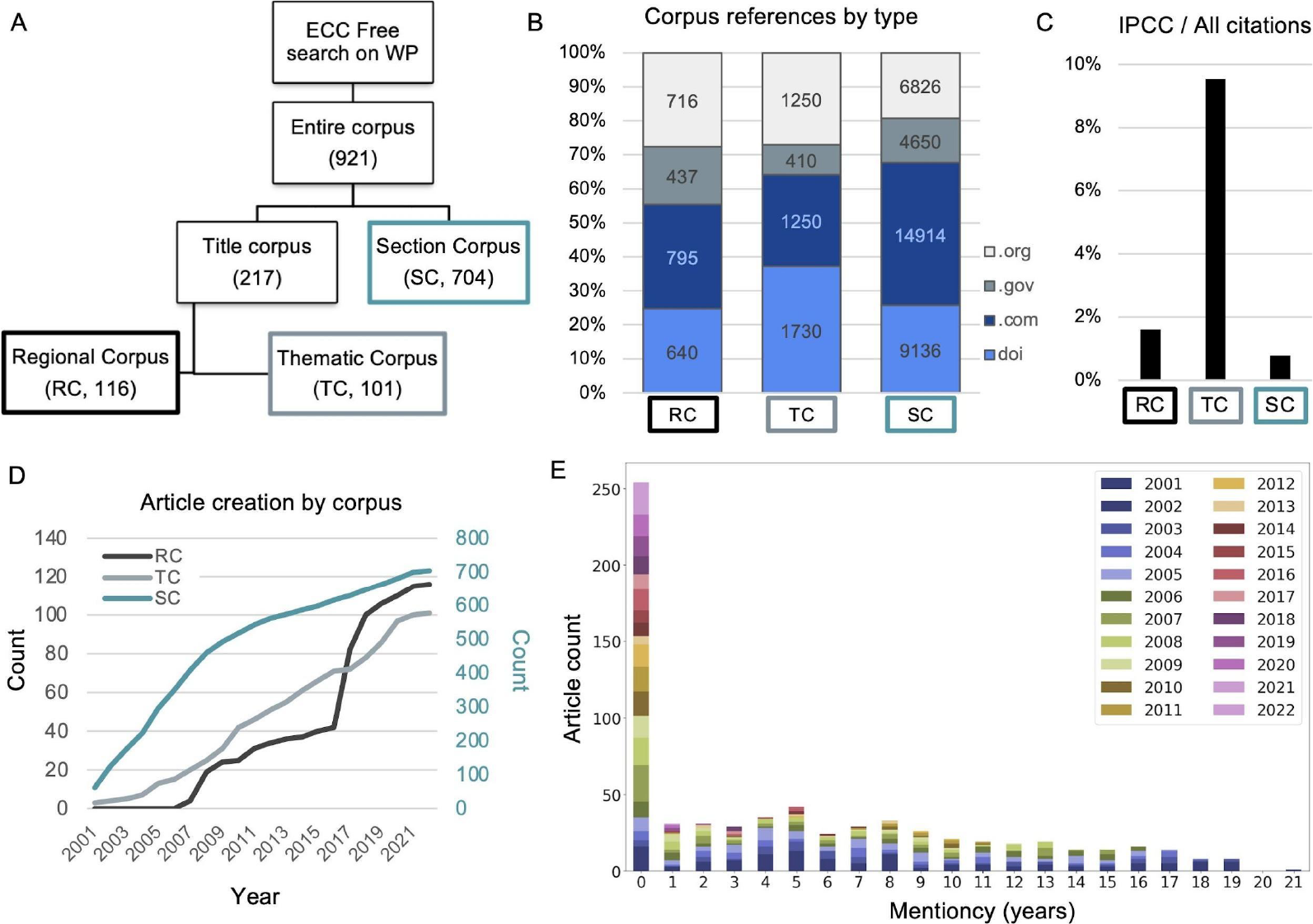
Effects of climate change on Wikipedia. A) Scheme of the corpus creation process: A free search of Wikipedia’s English-language articles on “Effects of climate change” to identify all relevant articles. These were then filtered to include only those with the term in either their title or that of a section (leaving 921 articles). The title articles were then filtered with the phrase “Climate change in” to create the Regional Corpus (RC) and the Thematic Corpus (TC). In brackets are article counts of each corpus. B) References distribution across the three corpuses. The numbers inside the bars indicate absolute citation counts, per type. C) The fraction of IPCC references among all references in each corpus. D) Timeline of the article counts per corpus since Wikipedia’s launch (2001). The articles titles and creation date (DOB) can be found in Table S1. E) Distribution of “mentioncy”: the duration of time between when an article was opened to when it first mentioned the term “climate change”. The bars are color coded based on the DOB. See also Table S1 and S3.

The “title corpus”, with 217 articles, was subsequently divided thematically, too; into a “Regional Corpus” (RC) that included articles on local effects of climate change (e.g., “Climate change in Antarctica”, or “Climate change in Scotland’’) and a “Thematic Corpus” (TC), for those with a wide perspective on climate change (e.g., “Effects of climate change on agriculture”). The “Section Corpus” (SC) contained the majority of articles (704) and touched on climate change-related topics (e.g., “IPCC Second Assessment Report”, “Environmental issues in Africa”) as well as other bodies of knowledge (e.g., “Polar bear” or “Ocean”).

The disparity between the corpuses show that climate change is both a field in its own right and one that has permeated into other bodies of knowledge, including those with more social or less scientific focuses (like “Fishery” or musician “Thom Yorke”). This divide, as we shall see, has to do with the growth not just of knowledge inside the field of climatology but also from a growing understanding of climate change’s widespread effects.

### 2.2 Corpus references and bibliometrics

Wikipedia’s guidelines demand all factual claims to be supported with a reliable source that can be verified by others. Sources and references are used not just within an article’s text to back specific claims, but also by editors themselves in justification of their edits (29).

We thus hypothesized these can be used to characterize the different corpuses. A bibliometric analysis revealed articles were supported by different types of sources: academic (containing “doi/PMID/PMC”), public (“.com” websites) and official (“.org” or “.gov”), (Fig 1B and Table S1). The distribution varied to some degree between corpuses: TC relied predominantly on academic sources, citing climate science. On the other hand, SC, in which climate science was nested within broader topics, had the majority of footnotes from public websites, such as news sites. This was also true but to a lesser extent in RC.

Among all these types of references we found a heavy reliance on IPCC reports, in line with previous studies on English Wikipedia (17) and interestingly also on the Portuguese version of the encyclopedia as well (18,19). However, IPCC reports did not always fall neatly into one type of bibliometric category, and were cited in a number of ways: At times studies published directly in IPCC reports were referenced through their DOI, while at other times interviews with researchers summing up their findings in news media outlets were referenced (.com). Therefore we added a specific IPCC count across corpuses and found TC had the highest prevalence of IPCC references, with over 9% of its references being that of IPCC reports (Fig 1C).

Contrasting the bibliometrics of the ECC corpus to another field of knowledge we recently researched using the same methods (9) allowed for comparison and contextualization of these findings. Comparing the overall percentage of academic sources to non-academic sources, a metric called “SciScore”, for the entire ECC corpus and a corpus of articles on CRISPR showed that climate change articles used less peer-reviewed studies than CRISPR articles (Fig S1A and Table S2). Gene editing was more scientific than ECC, at least on Wikipedia. Perhaps unsurprisingly, the CRISPR corpus made no use of IPCC reports as references.

These results show that different corpus of Wikipedia articles’ overall character can be reflected by the type of sources it utilizes, reflecting a connection between encyclopedic framing of content and bibliometrics.

### 2.3. Corpus historical dynamics

Wikipedia users can change the content of existing articles, as well as create new ones. The creation of a new article on Wikipedia indicates that a certain topic has reached a certain threshold of notability editors require for opening a new entry (30,31). Therefore, we hypothesized that tracking article creation can reflect the growing body of knowledge related to climate change that had accumulated between 2001 (Wikipedia’s creation) and 2022. To follow these dynamics, we used the data supplied by the “corpus builder” (9,28) to create a timeline of corpus articles based on each of their Date Of Birth (DOB), (Fig 1D and Table S1).

We found that as the corpus’ size (i.e. number of articles) increased over time, each subcorpuses’ growth followed a unique pattern. TC exhibited a relatively constant growth slope in terms of number of new articles. These include articles with a general climate-related theme (“Climate change denial”, 2015; or “Climate change and birds”, 2020) and those sparked by specific events (“2018 United Nations Climate Change Conference”, 2018) or the COVID-19 pandemic (“Climate change and infectious diseases”, 2020). RC articles began to appear later, only from 2007-2008, and then rose slowly until another spurt of new articles in 2018, perhaps indicating increased attention to specific locations and new bodies of knowledge addressing climate’s local ramifications.

Again, comparing the EEC corpus’ growth to that of CRISPR shows the two fields also display distinct growth patterns over time - much like the subcorpuses. CRISPR, an emerging field, peaks in terms of new articles at a much later stage in its history compared to ECC, whose major growth happened earlier (Fig S1B and S1C).

This analysis suggests the historical dynamics of article creation can serve to qualify the differences between fields and the manner scientific topics are represented on Wikipedia.

### 2.4. Cross-topic intersection

As noted, the majority of corpus articles were seemingly not directly linked to climate change, but rather only included knowledge from it as either a section or subsection (SC). To better understand the intersection of climate change with other bodies of knowledge we created a metric termed “mentioncy” - which gauges the time that passed between an article’s creation (DOB) and its first mention of the term “climate change” (Fig 1E and Table S3). We developed a freely available tool to automatically do this with any term - based on the already existing tool “wikiblame” (32), which searches an article’s edit history and compares past versions in search of the requested expression.

The mentioncy tool addresses questions regarding the temporal aspect of the topics convergence: were climate change and certain SC articles’ always connected or were they independent fields that only recently crossed paths? We found that nearly a third of the corpus articles mentioned “climate change” early - e.g., “Denialism” (from 2006). Among the >250 articles that included the term from the day they were opened are those on IPCC reports, weather-related events (e.g., “2016 Pakistan floods”, or “2020 California wildfires”) and many climate related articles like “Extreme Weather” or “Climate psychology”.

Other articles did not note the term upon opening, and their content could exist without requiring it editorially. Tellingly, the article for “Mindanao”, the second largest island in the Philippines, was the last to include the term climate change: it did so some 21 years after it was opened, in 2022, in wake of severe weather events that occurred that year. Similarly, the article for “Mindanao current”, an ocean current adjacent to the islands, mentioned climate change in 2021 (a 12 year lag, as it was created later) following the publication of a scientific paper in that year, which modeled climatic changes on ocean currents. These examples serve to highlight how changes in the world - be it real climatological shifts or the publication of knowledge about them - influences Wikipedia and spur cross-pollination of ECC with different fields.

## 3. A history of “Effects of Climate Change”

Our different corpus analyses showed Wikipedia can be used to follow the growing body of climate change related knowledge. To investigate climate change and Wikipedia with more historical nuance we decided to zoom in qualitatively on the article for ECC and use it as our “anchor article” - the main focus of textual historical analysis whose versions would be compared. This article was chosen based on its title and its representation of not just the field of climate (e.g., causes and manifestations), but its actual effects (Fig 2A).

**Fig 2:**
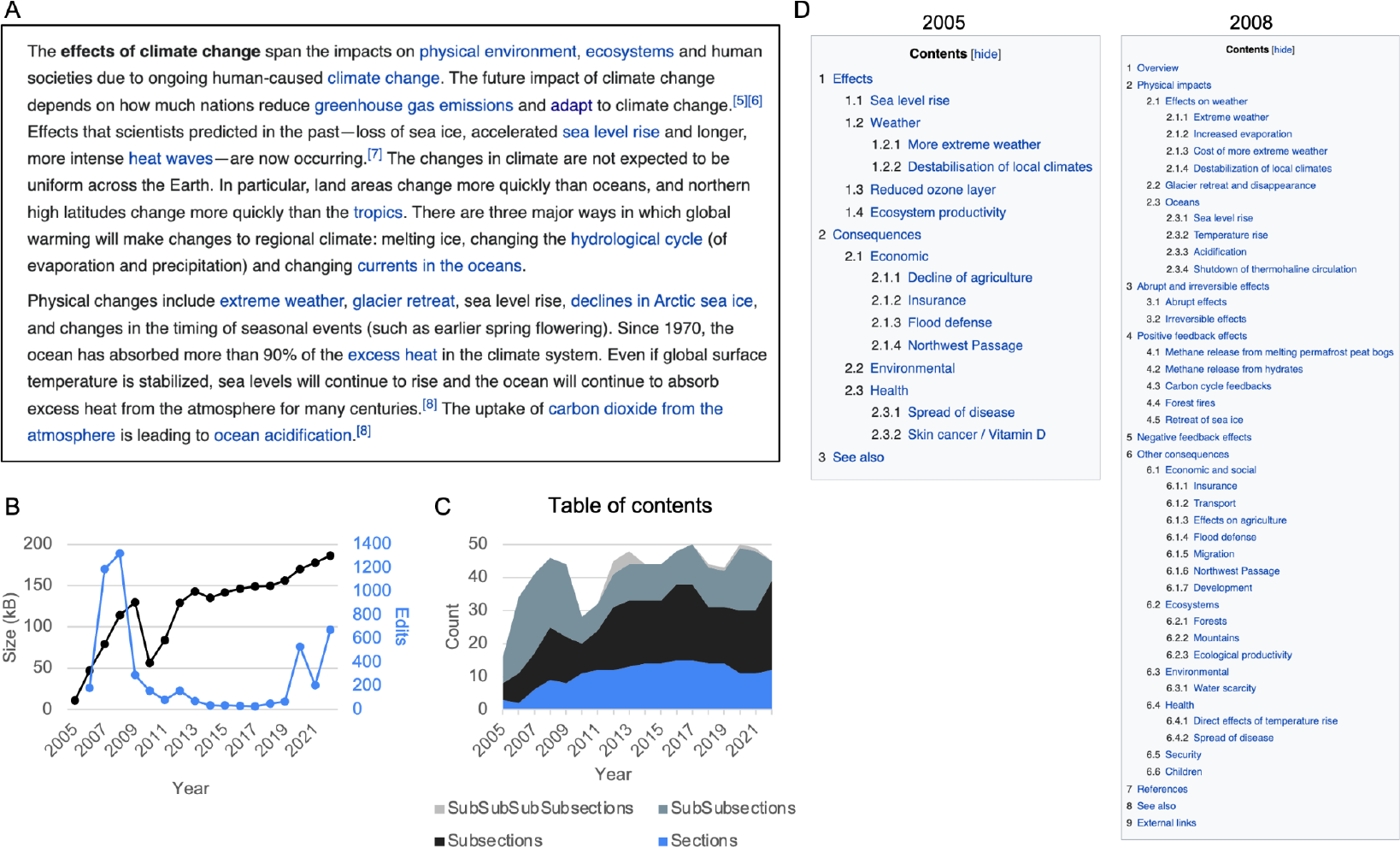
ECC anchor article. A) Lead of the ECC article on May 11th 2022. B) Article size and number of edits per year. C) Number of sections and subsections. D) Table of Contents (TOC) of selected years. More details of the TOC can be found in Table S4.

For this aim, we conducted a textual analysis of the article’s history, sampling it at annual intervals. Starting from its creation on June 26th 2005 until May 11th 2022, the text of each subsequent edition was compared to the previous one in terms of its lead section, Table Of Contents (TOC), references and key editors. This is a form of mixed methods meshing quantitative readouts and qualitative work (33), which can be labeled a form of “thick big data” (34) – a qualitative method involving a close reading and detailed review (35) of different datasets and their contextualization through “thick description” (36). In this case, our corpus served as our initial dataset, and it was contextualized by a close reading of textual changes to the historical versions of the article, with the anchor article’s edit history serving as a textual database and its metadata supporting the work.

Reading through its edit history, we found the text itself was a rich source of information about how scientific consensus about climate change and its consequences was displayed on Wikipedia. For example, in its first versions, the lead section of the ECC article addressed “the **predicted** effects of global warming… for the environment and for human life” (bold added). In the 2008 version, political or social claims regarding human responsibility to mitigate the phenomenon appeared: “Concerns have led to political activism advocating proposals to mitigate, eliminate, or adapt to it”, the lead section stated, now less hypothetical. Its latest version (from 2022) details the actual “effects of climate change [that] span the impacts on physical environment, ecosystems and human societies due to ongoing human-caused climate change” (Fig 2A). The shift between these different wordings took place in a gradual way. For example, the early text used the word “will” to describe the expected effects, while the more current versions use a past or present tense (“is warming”). These changes are in line with previous research showing the acceptance of the CC phenomenon as consensus (17). However, this historical process is not only reflected in the article’s text, but also in its form, its TOC and references, which can be both read and quantified. Therefore, the next sub-chapters present a quantitative analysis of these followed by qualitative ones, interspersing metrics with substantive examples gleaned from the article’s past versions to provide context and detail, in line with the tradition of thick description.

### 3.1. Structural changes

The TOC easily lends itself to mixed-method analysis, containing both semantic information gleaned from its text (the titles of sections and subsections) and quantitative metrics (e.g., number of sections or subsections over time). From its creation in 2005 until 2009 the article grew in both text size and form (TOC), (Fig 2B and 2C). In this early phase, the article’s content was more focused on predictions of future effects, like noted in the lead. Between 2009-2010, a dramatic drop in content occurred as part of intensive restructuring. From 2010 to 2014, both text and table of contents expanded again, staying relatively stable until the endpoint of this research (2022).

Despite high numbers in the article’s early years, size and edits were overall not highly correlated. They were however indicative of wider changes taking place in the table of content. Sections and their growth have previously been used by us (9) and others to map the growth of knowledge (37) on other scientific articles on Wikipedia. Alongside textual changes reflected in the article’s text and size, we also observed changes in its TOC, which expanded and contracted over time (Fig 2C and Table S4). From 2005-2007, the article’s sections were elaborated with subsections, for example, effects on “Weather” was elaborated to include the potential harm to “Oceans” (Fig 2D). The article retained its high number of sections (ca. 50) until 2022, although their content underwent restructuring, as indicated in shifts in the relation between sections and subsections. These shifts, we shall see, are linked to wider changes in Wikipedia’s climate articles and a broader reorganization or consolidation of its coverage of climate change’s effects overall.

Reviewing the qualitative shifts in the sections, their titles, and their order, revealed the nuanced dynamics laid out in broad strokes in the different historical phases. In its first years, the content of the article was broadly divided into two parts: “Effects” of climate change on the environment and nature (e.g., “Sea level rise” subsection), and its “Consequences” on various fields of human activities (e.g., “Economic” consequences like “Decline in agriculture” and others on human “Health”). Within this primary structure, the article continuously expanded with more scientific explanations for new observed climatic changes, alongside speculations about potential future ramifications.

An “Overview” section was created in 2006, discussing climate change causes, importance and looming impacts. Over the years “Effects” and “Consequences” still served as the main driving logic for the creation of new sections and then their expansion into additional subsections. For example, the subsection “Acidification” was added to the one focused on “Oceans” in 2006, and “Mountains” was added to the section on environmental consequences in 2007 (Fig 2D).

The addition of new effects in the form of new subsections continued until 2008, from which point they began to decrease: the result of some sections merging or forking off to form new articles, or others being removed. For example the content of section “Abrupt and irreversible effects” was moved into one titled “Destabilization of local climates” that dealt with melting ice. This subtle drop in number of sections foreshadowed the substantial reorganization that would take place between 2009 and 2010. The years 2009-10 indeed saw dramatic changes to the article, with both its structure and its content radically shifted. The modifications, we saw, consisted of summarizing the existing content: deletion, relocation and even forking off of entire sections to new articles. For example, the “Positive feedback effects” and “Negative feedback effects” sections were migrated to the article for Climate change feedback (opened in 2010).

Furthermore, the drop in ECC’s table of contents (Fig 2C) coincided with the small peak of articles created in TC (Fig 1D). We found, for example, that the “Increased evaporation” subsection was migrated to form the basis for the “Physical impacts of climate change” article by an editor called Enescot, who also created the article “Climate change, industry and society” from many of the subsections that were initially under “Economic and social” in ECC. Meanwhile, alongside these new articles, sections dealing with hyper-focused regional effects were removed, to the benefits of more broad explanations on chemical or geological mechanisms in play in the wider phenomenon of climate change. In 2012, alongside the growth in size, the article saw an increase in sections and subsections. Following this rise the TOC remains relatively static in numbers. “Benefits of global warming” was added and removed, short-lived throughout 2016-2017.

Editors who made these changes, as well as the 2010 restructuring, frequently justified their edits on the talk page (a forum-like arena where editors can discuss an article’s general direction or debate specific wordings or sources) by calling upon the publication of IPCC reports, which we have seen are highly cited in our corpus. The role of academic studies, as well as IPCC reports, will become clearer when contextualized within a wider analysis of the ECC’s references, the focus of the following chapter.

### 3.2. ECC references and bibliometrics

Next, we examined the sources used to support the information in the ECC article. The most cited references were IPCC reports, followed by academic papers, while websites or official sources were slightly less common (Fig 3A and Table S5). Delving deeper into the academic sources showed they come both from the field of climatology (e.g., Nature Climate Change) and from prominent general interest scientific journals (e.g., PNAS and Science), (Fig 3B).

**Fig 3:**
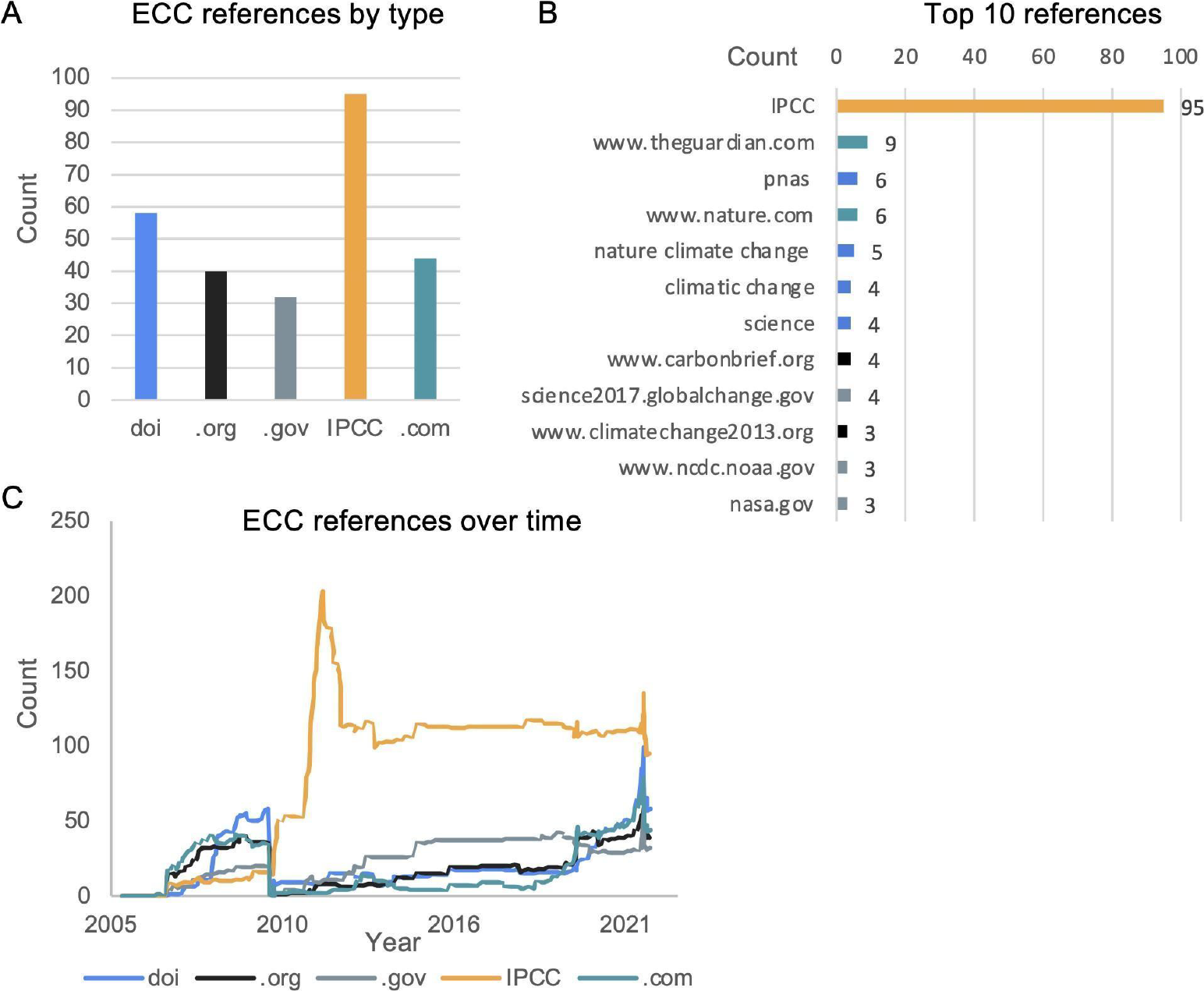
References and bibliometrics of the ECC article. A) References count, by type. B) Top 10 references in ECC, by number of citations. C) Historical distribution of references, by type. For more information see: Tables S5 and S6.

To understand the historical dynamics we analyzed these sources over time and tried to contextualize them within the edit history (Fig 3C and Table S6). Early versions of the article contained general predictions on possible consequences of sustained climate change, reflected in their sources: e.g., expected rise in ocean’s pH levels was added in 2007 and referenced the IPCC’s Summary for Policy Makers published that same year.

Following 2007, the IPCC reports became instrumental in the article’s ability to articulate the actual threat climate change’s effects increasingly posed. We found that all main sections had at least one IPCC reference by 2010. The section on sea level rising was completely revised that year and all its citations changed to IPCC references. A new section for “Specific health impacts” cited it almost exclusively, too, using it to back key paragraphs like “Food supply” and another in 2011 on food security. It is worth noting that the expression “level of confidence”, IPCC’s key term for expressing scientific certainty about climate change phenomena, was added to the ECC article for the first time in May 2010.

This trend continued and in 2011, at the height of the IPCC’s enrichment in the ECC article, there were only three citations out of 33 that do not refer to the 8th chapter of the 2007 report. This chapter focuses on the effects climate can have on human health, especially the increasing vulnerability to some conditions caused by the likes of storms, floods or diseases. In subsequent years the section titled “Health” significantly grew as new subsections and descriptions were added (Fig 2C).

The drop in article size, resulting from the aforementioned migrations, also coincided with a drop in references that year (Fig 3C) - but the IPCC remained highly cited. The IPCC’s influence on the article was even noticeable in its illustrations - with reports providing a source of copyright free figures as well as inspiration and data for infographics.

The findings of the 5th IPCC report (completed in 2014) were quickly integrated into the article following their publication. For instance, the footnotes section was expanded to explain key IPCC terms - such as the aforementioned “confidence levels” attributed to different predictions and potential climate scenarios. These were also linked to structural changes: in 2014, a section about scientific opinion also entered the article, and was (accordingly) based on academic references.

In 2019, as part of a Wikipedia-wide initiative to reformat footnotes, we observed a decline in official sources. While IPCC sources formed 50% of the article’s references in 2019, they were down to 39% by May 2022 (Fig 3C). However, the drop in absolute number of IPCC references was in fact minor and resulted from the addition of new references from academic journals and newspapers like *The Guardian*, which had disappeared from the article in 2010, but was reintroduced in 2019.

### 3.3. Editors

To gain insight into the editorial considerations supporting the addition of this content we also examined those behind it, going backstage to scrape user data from the ECC article. Here our mixed methods included quantitative edit counts over time and qualitative thick description of editorial activities, gleaned through the history log and talk pages.

Interestingly, a large proportion of the article’s edits were made by only a handful of users (Fig 4A). Indeed, the top three editors are responsible for nearly half of the edits made to the article from its creation, a small number compared to the fact that it has been viewed over 4 million times since it was opened (38).

**Fig 4:**
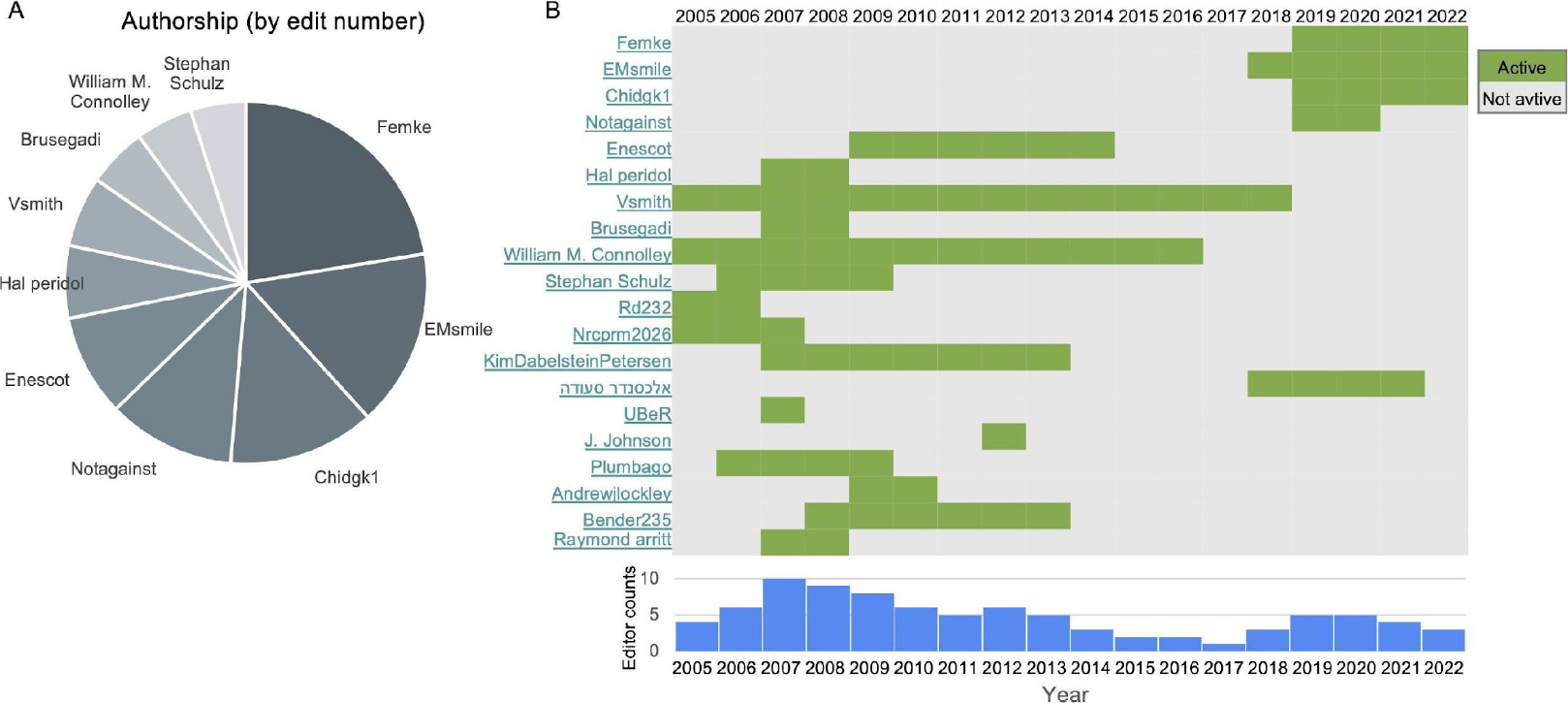
Prominent users in the ECC article. A) Top 10 editors, based on edit count. B) User activity timeline of the top 20 users. In green are years of activity for each user. On the bottom are counts of active users per year (out of these 20).

From the time they first opened the article in June 2005 and until roughly the end of September that year, the user Rd232 was the main editor of the article (Fig 4B). They set the initial tone and framing, creating the article from content they migrated from the article on global warming. However, from then on, Rd232 seems to have retired from his editorial activities, and other editors emerged in his stead. Most of the 2010 restructuring was led by a single user - Enescot. The IPCC, specifically Confalonieri et al., was featured prominently in the post-2011 period in wake of this editor’s work, which went beyond adding just references and also included editing content, reworking sections, as well creating new articles based on sections they migrated from ECC.

The Wikipedia user Femke first edited in August 2019 and soon after became the top editor of this article. They were also involved in a series of arguments on ECC’s talk page. Femke was worried about the quality of sources, and after lamenting the issue on the talk page began to replace outdated sources, reports and data in the article’s text. Among other things, they removed a section on air pollution and also replaced old emission scenarios laid out by the IPCC in 2000 with more recent ones (from 2019 and 2021).

Breaking-down the qualitative content of the editors’ work highlights two things. First, IPCC reports played a distinct and unique role in making scientific knowledge on climate change accessible as a source. Second, the high impact that body of references had was the result of the work of specific editors who undertook upon themselves the task of restructuring and then expanding the article based overwhelmingly on IPCC reports. The major role some editors may play in shaping an article has interestingly been highlighted in climate change-related articles on Portuguese Wikipedia (19), but also showcases how freely accessible scientific knowledge can find its way to the public thanks to the volunteer work of a handful of people.

## 4. Discussion

In summary, we found that Wikipedia’s articles documented the discourse around effects of climate change for over 20 years, in terms of the number of articles dedicated to what was initially treated as a hypothesis, and then in terms of scope as the expected effects became a forlorn reality. Alongside ECC and its corpus’ growth, we also saw knowledge from this space penetrate other fields over time. The main article on ECC changed and, much like the articles that grew from it, made much use of IPCC reports in its text, structure and references. The IPCC had a big impact on Wikipedia and its history, we found, and what made this possible was the work of a small number of dedicated editors.

In recent years, Wikipedia has been increasingly established as a historiographical source (3,8,15,37). Previous work regarding the Wiki-article for “Global warming” between 2002-2014 concluded that the IPCC report was prominently used (17). Furthermore, they found that IPCC’s usage was, at least initially controversial, and that alike the controversiality surrounding CC in general, subsided in the period following 2010-4, reaching consensus status. Our results reinforce their findings and elaborate on them: After the 2010 restructuring IPCC remained the main reference of ECC article, while being frequently used across the whole corpus (Fig 1B and 1C). After inevitably becoming an unavoidable argument on the talk page, more and more claims were made supported by one or the other of the IPCC reports, which thus became what could be coined an “obligatory passage point” (19,39) for any editor willing to contribute to the article productively and counter denialism with science.

This IPCC’s prominence highlights how Wikipedia remains in lock step with science in terms of societal attempts at combating denialism. For example, alongside the widespread use of highly rigorous and well-respected climate studies that were well inside the scientific consensus, the term climate change entered the article “Denialism” early (2006) and another article dedicated to climate change denial which was created in 2007.

Wikipedia also documented these debates: another major historical change recorded by the article’s text was its title: In 2020, the article was renamed “Effects of Climate Change” after almost a decade and a half as “Effects of Global Warming”. This was not just true for our anchor article: many articles initially included the term global warming but were rephrased - even the main article “Climate Change” was first dubbed “Global Warming”. This followed a much wider process playing out outside of Wikipedia and backed by science and policy makers, too, to shift public discourse from global warming to climate change (40). In 2005, the National Academy of Sciences published a pamphlet to explain that “climate change” is a scientifically more relevant framework than “global warming” (41). The following year the U.S. Environmental Protection Agency changed its website domain from global warming to climate change (42). Over the past 20 years, the terminology has shifted and led to a gradual adoption of the term climate change over global warming, a process documented anecdotally on Google Trends (43), social media (44) and media outlets (45). This transition stems from the different implications of the two terms: global warming relates to the trend of increasing average global temperatures; alternatively, climate change is a much broader term, including alterations in climate patterns, including but not restrictive to warming, for instance increased levels of atmospheric carbon dioxide resulting from fossil fuel use. On Wikipedia this process was documented first over years of debates on articles like ECC, and eventually with the title shifts and its implementation across the English-language encyclopedia at the corpus level.

Alongside the historical value of its text, title, and edit history, Wikipedia’s references provide a rich source of bibliometric data with historical significance, too. Bibliometrics is a well established practice within the history of science (46) and has long been used in historical research (47,48) and digital humanities (49). Past studies into the bibliometrics of scientific topics on Wikipedia found that these are linked to wider trends in scientific knowledge (3,9), and tend to be from high-impact factor journals from both niche journals or general ones (2,11). In the case of the ECC and our corpus, this was also found to be true: we saw that among those academic journals cited there were prestigious field-specific journals like Nature Climate Change. While research has shown that scientific articles predominantly use academic sources, or at least had a clear bias towards them, ECC use of IPCC reports well went beyond just academic sources. The reports were cited in numerous ways, showing how domain-specific yet also high-impact research can be referenced in Wikipedia in a way that challenges the division between academic, official and public sources (DOIs, .com, .org). For example, alongside different types of IPCC references, we could also see the use of official data linked to national and local governments cited in articles about regional effects of climate change (.gov). This joins a number of well documented technical issues and methodological challenges that conducting bibliometrics analyses on Wikipedia face (50). More importantly, as we and others (17) saw, the IPCC’s effect was not limited to bibliometrics alone: IPCC reports were used as references but they also informed the ECC article’s content, structure as well as the corpus itself - with new articles forking off from the main article, all to conform with IPCC thinking over time.

Our findings highlight the unique role IPCC reports played in making scientific knowledge about climate change actionable for the public, and underscores Wikipedia’s ability to facilitate access to it. The fact a single source exerted such widespread effect on the articles, was also partially the result of the work of single editors exerting their influence and enforcing a rigid academic sourcing standard. This dominance of a handful of Wikipedians on one specific article or topic’s editorial history is in fact not uncommon (51). The article’s original framing and form was laid by Rd232, the editor who opened it and thus defined its initial trajectory (52). As the Nobel Peace Prize was awarded to the IPCC and former US Vice President Al Gore, the year 2007 was a significant one in the field, which attracted much attention to the ECC topic, which is likely reflected in the article’s spike in editorial activity around this time. Most of the 2010 restructure was led by a single user, one of the top ten contributors on the article, and included the addition of the Confalonieri et al. reference, which was the 8th chapter of the 2007 report, and was featured prominently in the post-2011 versions. The editor who added the IPCC reference and led the structural reorganization in its wake, Enescot. They explained the logic for using the IPCC source in an editorial discussion held on the article’s talk page, saying it “went through extensive review and has been accepted by most of the world’s governments [and party to the United Nations Framework Convention on Climate Change] as providing a ‘comprehensive, objective and balanced view of the subject matter.’” Their use of the source in the text, not just as a reference but also as a structural force was, in their own words, due to the rigid scientific review process they underwent and their status within the scientific community. The fact that dedicated editors can play such a massive role in facilitating access to scientific knowledge and translating research into the public discourse has also been anecdotally seen in other fields. For example, the field of sleep and circadian clocks on Wikipedia were overseen by a single editor suffering from a severe sleep disorder (3).

This study compared three corpora of hundreds of articles dealing with climate change. It further compared them to a CRISPR corpus, thus allowing us to understand how contemporary science manifests on one of the world’s top websites. Together these comparative findings and the tools behind them show how researchers can use Wikipedia as a trove of open data and knowledge to enrich our understanding of the growth of knowledge and its representation online. Our findings are in line with previous works conducted on Wikipedia and climate change (17–19) as well as other scientific fields with similar methods (9,12). This work, which we associate as a form of “thick big data”, helps underscore how computational and qualitative analyses are not just possible but lucrative, and help go beyond a divide that has long plagued research on Wikipedia. These shared methods, based on the open tools available to others to use freely, result in comparable works on different fields on Wikipedia (53). Together, a complex and eventful history of the most controversial and important scientific issues of our age, can be created.

## Code accessibility

Our code can be found at: https://github.com/RonaTheBrave

## Acknowledgements

The authors wholeheartedly thank Ariel B. Lindner for his support and valuable guidance, as well as Jean-Marc Sevin and Leo Blondel for their computational groundwork. Thanks to the Bettencourt Schueller Foundation long term partnership, this work was partly supported by the LPI Research Fellowship, Université de Paris, INSERM U1284, to RA and OB. RA’s work was supported in part at the Technion by a fellowship of “The Israel Academy of Science and Humanities”. In either case, the funders had no role in study design, data collection and analysis, decision to publish, or preparation of the manuscript.

## Supporting information

**Fig. S1:**
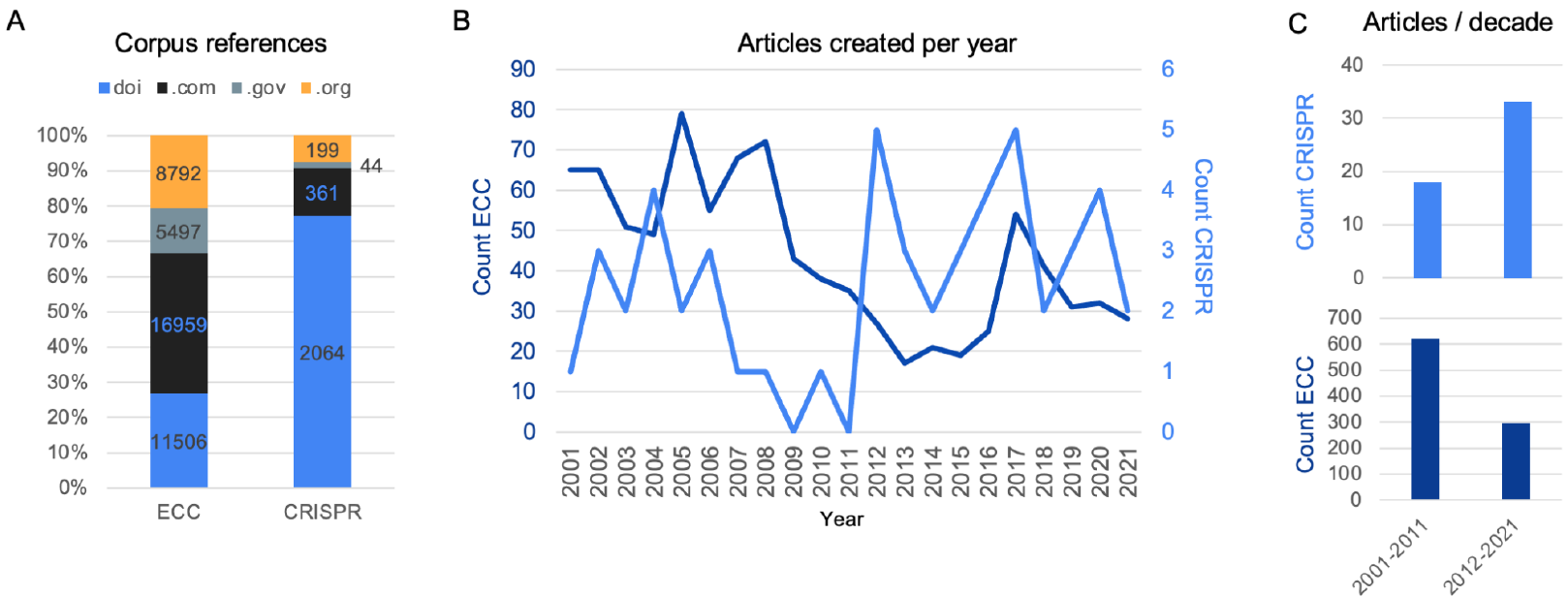
Comparison between ECC and another field - CRISPR. A) Sources distribution across different types, comparing the entire ECC corpus (921 articles) and the CRISPR corpus (51 articles). The bar plot presents the ratio of the sources and the numbers indicate the absolute counts, per type. B) Number of articles opened, per year, per corpus: ECC (dark blue) and CRISPR (light blue). C) Sum of B, per decade. CRISPR analysis was performed in February 2022.

## Supporting information: Tables

Supplementary tables are available online: https://sandbox.zenodo.org/record/1226798

